# Influence of Housing, Sex, and Sampling Location on Taxonomy and Function of Adult Zebrafish Microbiomes

**DOI:** 10.1101/2025.05.08.652836

**Authors:** Jeremiah W. Wanyama, Lydia Okyere, Luoyan Duan, Christopher A. Gaulke

## Abstract

The zebrafish (*Danio rerio*) has emerged as an important animal model for the study of host-microbiome interactions. However, information on how variation in experimental parameters contribute to microbiome structure and function in adult zebrafish is limited which complicates experimental design, interpretation of results, and may reduce reproducibility. Here we quantified the impact of two potential sources of microbiome variation – housing strategy and sampling location – on microbial diversity of adult zebrafish using 16S rRNA amplicon sequencing. Our findings indicate that housing strategy significantly impacts gut microbiome diversity in adult fish with the highest similarity between individuals co-housed on recirculating water systems. Microbiome acclimation after housing transfer took between 14- and 21-days. Significant variation in microbiome composition was also observed across sampling sites. As in humans, fecal and intestinal microbial communities were similar and varied by sex, however each body site sampled possessed a small site-specific microbial community signature. Consistently, imputed function of these communities showed that gene family diversity is also predicted to vary by body site particularly between gut and non-gut locations. Together our work demonstrates that housing, sex, and sampling strategy all significantly impact microbial community composition and highlight the need for community wide discussions on best practices and reporting standards for adult zebrafish microbiome studies.

## Introduction

Over the past four decades, use of laboratory zebrafish (*Danio rerio*) has grown from a niche experimental system for developmental biology to a critical animal model supporting innovation and discovery in dozens of fields ^1–4^. The adoption of the zebrafish has been spurred by its low cost, high fecundity, ease of genetic manipulation, and genetic and physiological similarity to humans^5^. Recently, the zebrafish has emerged as a promising model of host-microbiome interactions and has been used to demonstrate translationally relevant causal relationships between microbiome function and host health^4^. The adaptability, high-throughput, and extensive suite of molecular and genetic tools mean that this model system has the potential to massively accelerate the pace of mechanistic discovery in microbiome research ^4,6^. However, limited information on how choice of experimental parameters impacts variability in outcomes constrains utility of this model and threatens the reproducibility of zebrafish microbiome research.

Extensive work demonstrates that factors such as diet, sex, environment, and sampling location significantly impact microbiome composition and can affect interpretations and reproducibility of experiments^7–12^. To mitigate these impacts, best practices and reporting standards for microbiome research have been developed for mice and humans and have helped guide researchers in experimental design^13–15^. Best practices for microbiome investigations in companion animals^16^, wildlife^17^, and gnotobiotic larval zebrafish have also been developed ^18,19^. Although general guidelines for the husbandry and welfare of zebrafish have been well established ^5,20^, beyond dietary impacts^21–24^ little is known about how major sources of variation in mammalian models influence microbiome experimental outcomes in adult zebrafish. Defining if and how these experimental parameters shape microbiome variation is a crucial first step in identifying best practices and standards for reporting for microbiome experiments in adult zebrafish. Since cataloguing all sources of variation will require considerable time, understanding how the principal drivers of microbiome variation in mammals^13–15^ impact zebrafish microbial communities should be prioritized. Amongst the most critical considerations for microbiome studies in preclinical animal models are housing and sampling strategy^14,15^.

Housing conditions and housing density are major determinant of microbiome variation and experimental outcomes in mice^7,13,14^. Co-housing is often desirable due to reduced cost, but can result in microbiome homogenization across animals due to proximity and coprophagia^13,14^. Individually housing animals can prevent homogenization, but may be stressful in social animals such as mice and zebrafish^25^. In zebrafish, individual housing allows longitudinal profiling of the microbiome in individual animals^26^, however its impacts on gut microbial communities are not fully understood. It is also unclear how long gut microbial communities may take to acclimate to changes in housing. Quantifying the impact of housing and time needed for acclimation would help ensure microbiome variation across experimental groups is attributable to experimental variation and not equilibration to new housing conditions.

Another major contributor to variation in microbiome research is sampling strategy^8^. In mammals, fecal sampling is a popular non-invasive strategy for microbiome profiling. A major limitation of this approach is that the composition and function of fecal microbial communities poorly reflects small intestinal and tissue associated microbiota^27–29^ making it a suboptimal choice for some research questions^28^. These differences in microbiome profiles are driven by distinct physiologically and anatomically conditions along gastrointestinal tract^27^. In adult zebrafish, whole intestinal and fecal sampling are the primary microbiome sampling strategies^21,22,26,30^. While zebrafish gut anatomy is less complex than that of humans, several distinct microenvironments exist across the length of the zebrafish intestine which manifest analogous physiology to human gut compartments^31–33^. However, it is unknown if these intestinal microenvironments host a distinct microbiota in adult zebrafish making it unclear if fecal sampling reflects the entire gut microbiome or just specific compartments ^30,34,35^.

Here we examined how housing and sampling strategy impacted microbial community diversity in adult zebrafish. To gain deeper insights into the potential functions these microbes may perform at different anatomical sites, we imputed the functional potential of these communities and examined associations between function and location. We find housing have substantial impact on microbial communities and that when system type or housing is switched, the gut microbiome undergoes rapid remodeling. Taxonomic and functional diversity also varied significantly across body sites and between sexes. Each location investigated possessed unique diversity, and similar body sites (*i.e.*, gut tissues) shared a substantial fraction of diversity compared to distal sites mirroring ecological compartmentalization seen in mammals^27^. Together our findings suggest that, as in mammalian microbiomes, housing, sampling strategy, and sex all significantly impact research findings and highlights the need for development of best practices in adult zebrafish microbiome research to ensure the reproducibility of findings across labs.

## Materials and Methods

### Zebrafish housing

Animals were housed on a temperature controlled (28°C) Aquaneering continuous recirculating flow or timed water exchange flow-through aquatic system with a 14h on 10h off light/dark cycle. Both aquatic systems were housed in the same room and received the same source water. Water quality was maintained using automated dosing systems at 7.0-7.5 pH, 500-1200 µS, and 0ppm NH_3_, 0ppm NO_2_^-^, 0ppm NO_3_^-^. Animals were housed at a density of five fish per liter or less (group; 2.8 L tanks) or individually (1 fish / tank; 1.8 L tanks) and were fed Gemma micro 300 (Skretting, Westbrook, ME USA) twice daily. A total of 54 adult T5D line zebrafish were used for this work (N = 27 for flow and housing experiments; N = 15 for sampling location experiments; and N = 12 for investigations of the impact of sex on the microbiome). Animals used for this work were 12-14 months old and each experiment sourced animals from a single distinct spawn. All experiments involving animals were approved by the University of Illinois at Urbana-Champaign Institutional Animal Care and Use Committee.

### Sample collection and processing

Fecal samples were collected from adult tropical 5D line (T5D) zebrafish as we have previously described^36^. For longitudinally sampled animals used to study the impacts of housing, after morning feeding, animals were transferred to individual 1.8L tanks and monitored until defecation. Fecal samples were collected using sterile transfer pipets into sterile nuclease free microcentrifuge tubes. Animals were returned to their original tanks and resampled at each subsequent time point until necropsy. All other animals were sampled once at terminal timepoints immediately before euthanasia. Animals were euthanized by rapid chilling in ice water bath immersion.

For a subset of animals (N=15) the operculum was resected to expose the gill which was gentle swabbed using a sterile cotton swab (Hardy diagnostics, Santa Maria, CA, USA) for 5 seconds. Skin swabs were collected from animals by gently swabbing the dorsal and ventral skin surfaces of the fish 3 times with a sterile cotton swab (Hardy diagnostic, Santa Maria, CA, USA). The gastrointestinal tract was then removed (esophagus to anus) and the remaining gut cleaned of adherent non-gut tissue. The gut was then cut into three segments approximately equal in length (proximal, middle and distal regions). The swim bladder was also collected and cryopreserved. All samples were collected in sterile nuclease free microcentrifuge tubes and stored at -80°C until further processing.

### DNA extraction, library preparation, and sequencing

Microbial DNA was extracted using microbial DNeasy Powersoil Pro Kit (QIAGEN, Germantown, MD, USA) according to manufacturer’s protocol. An additional ten-minute 65°C incubation was added to facilitate cellular lysis prior to bead beating (20m) using a Vortex Genie 2 (Scientific Industries, Bohemia, NY, USA). Purified microbial DNA samples were eluted and stored at -20°C. Amplicon libraries were prepared as described previously^37^. Briefly, the 16S rRNA gene was amplified in triplicate using barcoded sequencing primers directed against the V4 region (515F and 806R) ^38^. Triplicates PCR amplicons were pooled, visualized on 1% agarose gel to confirm expected fragment size, and concentrations quantified with the Qubit HS kit (Thermo Fisher Scientific, Waltham, MA, USA). A total of 200ng of each library was pooled together and cleaned using QIAquick PCR amplification Kit (QIAGEN, Germantown, MD, USA). The amplicon pool was then sequenced at the Roy J. Carver Biotechnology Center at University of Illinois at Urbana-Champaign (300bp PE) on an Illumina Miseq. The resulting paired end amplicon reads were quality controlled, denoised, and filtered of chimeric sequences (default parameters) using DADA2 (v1.20)^39^. Taxonomy was assigned to the remaining ASVs with DADA2 using the SILVA reference database (v138.1)^40^.

### Microbial diversity and differential abundance analysis

Amplicon sequence variant abundances were normalized by rarefaction or relative abundance (vegan v2.6.6.1) in R. Richness and Shannon entropy were calculated using vegan and differences between conditions were quantified using generalized linear mixed models with glmmTMB (v 1.1.9). Microbial beta-diversity was measured using Jaccard distances and ordinated with non-metric multidimensional scaling (NMDS). Associations between beta-diversity and experimental parameters was assessed using Permutational Multivariate Analysis of Variance (PERMANOVA; vegan::adonis2 v2.6.6.1, permutations = 5000). Differences in intra- and inter-individual beta-diversity (vegan::vegdist; method = “jaccard”) were quantified with two-tailed pairwise Wilcoxon rank sum tests (holm correction).

Negative binomial general linear mixed models (glmm; glmmTMB v1.1.9) were used to quantify differences in microbial abundances between groups. Briefly, for each taxon, a null model containing only the intercept was compared to an alternative model that contained intercept, tissue or intercept, tissue and sex as predictors. An ANOVA determined if the alternative model explained more variation than the null model. False discovery rate (FDR) was controlled with the Benjamini & Hochberg method (R stats::p.adjust, method = “BH”). This same strategy was used to evaluate the impacts of housing and aquatic system type taxa abundances. A FDR threshold of 0.1 was used unless otherwise indicated. The number of unique and shared microbial taxa were defined as taxa whose abundance met a specified prevalence threshold in only the defined subset of tissues. Shared taxa were visualized with the Upset R package (v1.4.0).

### Functional imputation analysis and differential abundance testing

Microbiome functions were imputed from rarefied sequence variant tables with PICRUSt2 (v2.5.2)^41^ using max parsimony. Associations between KEGG protein family diversity, pathway diversity, tissue location and sex were measured as above with a PERMANOVA (5,000 permutations). Kruskal-Wallis rank sum tests quantified differences in protein and pathway abundances and false discover rate was controlled at 0.005 using the Benjamini & Hochberg procedure.

KEGG pathways were quantified by summing the abundance of the KEGG orthologous gene families they contained. Functional categories were based on BRITE categories with some being combined into broader functional groups. For example, CALEN Metabolism contains the BRITE categories: amino acid metabolism, carbohydrate metabolism, lipid metabolism, metabolism of other amino acids, energy metabolism, and nucleotide metabolism. Metabolism of secondary metabolites contains the BRITE categories: biosynthesis of other secondary metabolites, metabolism of terpenoids and polyketides, and Metabolism of cofactors and vitamins. Drug resistance contains drug resistance: antimicrobial and drug resistance: antineoplastic categories. The other category contains the BRITE categories: global and overview maps, folding, sorting and degradation, membrane transport, translation, cell growth and death, nervous system, signal transduction, and transcription. In addition, the BRITE category Cellular community – prokaryotes was relabeled as Biofilms and quorum sensing since the only pathways of interest here were related to these functions.

### Data availability

All sequencing data used in this manuscript is deposited at the National Center for Biotechnology Information Sequence Read Archive under project number PRJNA1204388 samples SAMN46040644 - SAMN46040890.

## Results

### Housing type substantially impacts the gut microbiome

Husbandry conditions are known to strongly influence the composition and ecology of host associated microbiomes and considerable work has been focused on defining best practices for minimizing these effects in murine models^7,42,43^. However, in zebrafish, information on the impacts of husbandry parameters on the microbiome is limited. To begin to address this gap in knowledge, we asked if aquatic system flow type impacted the composition of adult zebrafish microbial communities. We focused on the two most widely used types of flow in aquatic systems: recirculating – water passes through tanks, is filtered, and passed through a high intensity ultra-violet light before passing back through tanks – and flow-through – conditioned water is passed through tanks and then directly into the drain. Adult T5D zebrafish (N = 27) were transferred from group housing on a recirculating flow system to group or individual housing on a flow through system. Animals were then allowed to equilibrate for 21 days with fecal samples collected every weekly (**Supplemental Figure 1A**). Fecal microbiome richness – the number of taxa in a community (**Figure 1A**) – and Shannon entropy – a measure of community randomness and richness (**Figure 1B**) – both rapidly and significantly decreased (glmm; richness: z = -6.01, *P* = 1.84x10^-9^; Shannon: z = -3.78, *P* = 1.57x10^-4^). Shannon entropy, but not richness, was also decreased for individually housed animals (z = -2.05, *P* = 4.02x10^-2^). Similarly, microbial community diversity (**Figure 1C**) associated with flow type (PERMANOVA; R^2^ = 0.15, *P* = 2.00x10^-4^), housing type (R^2^ = 0.06, *P* = 3.99x10^-4)^, day (R^2^ = 0.19, *P* = 2.00x10^-4)^, but not tank of origin (R^2^ = 0.02, *P* = 1.4x10^-1^). To evaluate how water flow and housing impacted specific taxa, we quantified differences in genera abundance using generalized linear mixed models. A total of 169 genera were significantly impacted by choice of system (FDR= 0.1) many of which are major components of zebrafish gut microbiota (**Supplemental Table 1**). For example, the genera *Aeromonas* (glmm; z = 7.29, *P* = 3.12x10^-13^) and ZOR0006 (z = 2.87, *P* = 4.11x10^-3^) were significantly increased and *Vibro* (z = -4.07, *P* = 4.78x10^-5^) and *Nitrospira* (z = -2.39, *P* = 1.70x10^-2^) decreased on the flow-through system (**Supplemental Figure 1B**). Housing also significantly (FDR = 0.1) impacted several highly abundant taxa with decreases in ZOR0006 (z = -4.77, *P* = 1.89x10^-6^) and increased *Aeromonas* (z = 5.63, *P* = 1.77x10^-8^) in individually housed animals (**Supplemental Table 2**).

**Figure 1.**
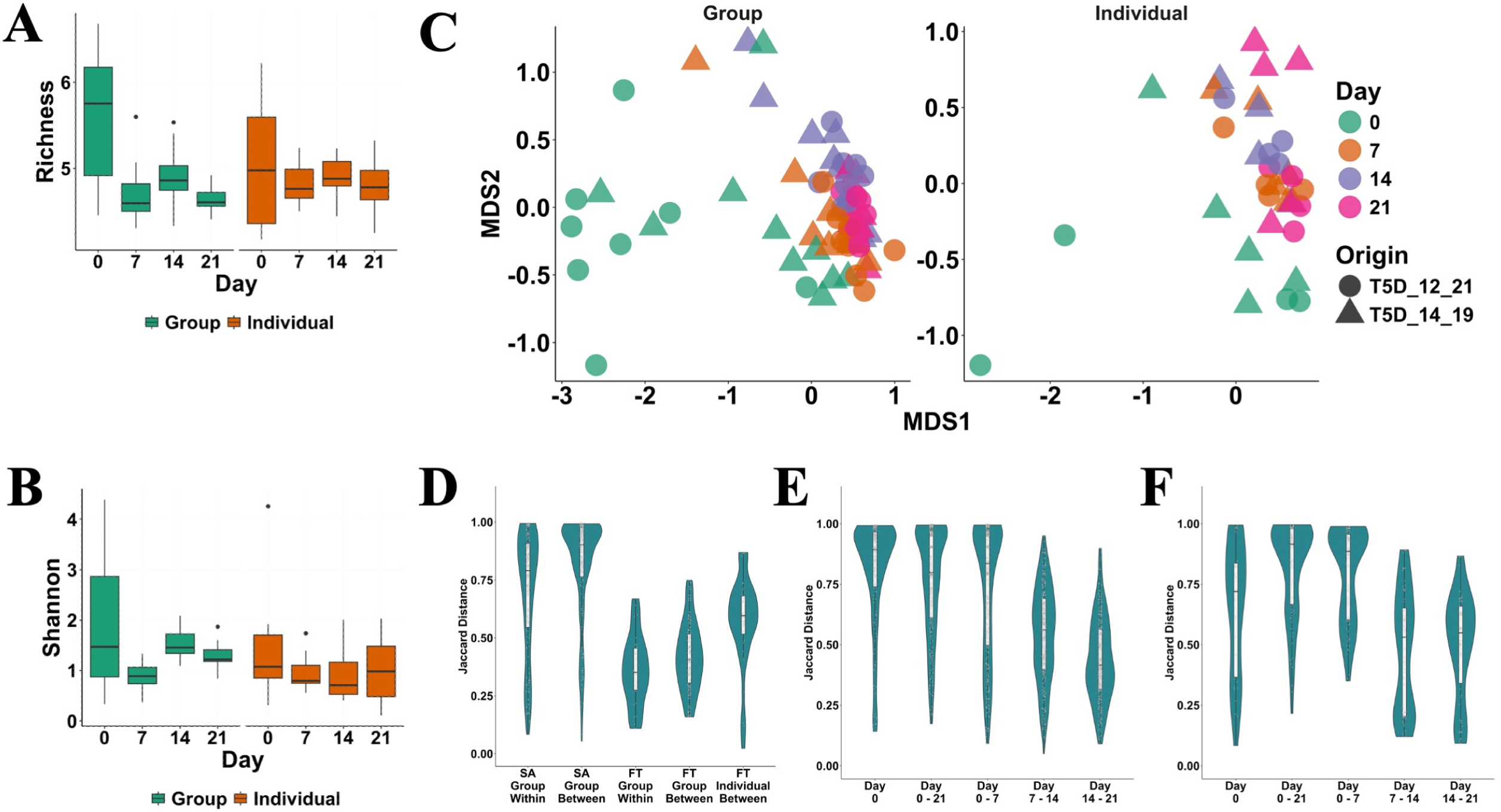
Husbandry substantially impacts zebrafish microbial community diversity. Box plots show distribution of amplicon sequence variant (ASV) **A)** richness and **B)** entropy in animals housed on a stand-alone (SA) recirculating flow rack (Day 0) or on a single pass flow-through (FT) zebrafish housing rack (Day 7-21). **C)** An ordination of non-metric multidimensional scaling of microbial communities from group (left panel) and individually (right panel) housed zebrafish across 21 days of equilibration to the FT system. **D)** Comparison of Jaccard distance (dissimilarity) between animals housed in the same tank (within) or in different tanks (between) tanks in group and individually housed animals. Time course analysis of Jaccard distance between equilibration time points in **E)** group, and **F)** individually housed animals.

In mouse models, animals housed together tend to possess microbiomes that are more similar to one another than to animals housed in other cages^14,43^. While it has been assumed that this phenomenon would manifest in zebrafish as well, it has not been extensively investigated. We found that, as in murine models, zebrafish housed together on recirculating flow systems (*i.e.*, housed in the same tank) possessed microbiomes that were more like each other (intra-individual variability) than animals housed in other tanks (inter-individual variability) (**Figure 1D**; *P* = 3.40x10^-5^). A similar trend was observed on flow-through systems, but it did not reach significance (*P* = 5.21x10^-2^). Overall, levels of intra-individual (*P* = 1.30x10^-8^) and inter-individual diversity (*P* < 2.00x10^-16^) were higher in group housed animals on recirculating systems. Individually housing zebrafish on the flow-through system increased inter-individual diversity relative to group housed animals (*P* = 5.40x10^-6^), however, variability was still lower than group housed animals on recirculating systems (*P* = 9.50x10^-9^).

Given the differences in microbial community diversity observed between flow-through and recirculating systems, we next asked how long it takes a microbiome to acclimate to new housing conditions. To do this, we first computed the Jaccard distances between microbiomes at successive time points, which represents how different animals are across time, in group (**Figure 1E**) and individually (**Figure 1F**) housed animals. Next, we computed Jaccard distances for each pair of animals at each time point, which represents how similar animal microbiomes are at a given time point. We considered the animals to be acclimated when the difference between successive time points was not statistically different from the distances between animals (*i.e.*, when temporal variation was approximately equal to population variation). As expected, the largest distances (lowest similarity) were observed at Day 0 (recirculating flow samples), day 0-21 (across the whole acclimation time frame) and day 0-7 (the first timepoint after transfer to the flow-through system). In individually housed animals, differences in temporal and population distances failed to reach significance at day 14 (Pairwise Wilcoxon, *P* = 1.00) while equilibration occurred later in group housed animals being delayed to day 21 (*P* = 0.78). Collectively these data indicate that animals housed on recirculating systems possessed microbial communities that are more individually distinct than those housed on flow-through systems, and that individual housing animals on flow-through systems somewhat reduces this effect.

### Zebrafish microbial diversity varies across sampling locations

In humans, differences in composition also manifest between gut tissues (e.g., stomach, large intestine and small intestine) and feces^37,44,45^ meaning that fecal samples may not accurately represent microbial communities at mucosal tissue sites. This is particularly problematic when disease specific patterns in the microbiome only occur in mucosal as sampling of feces exclusively can miss important biomarkers of disease^37^. In zebrafish, fecal and intestinal sampling are common, however, it is unclear if similar differences in microbiome across gut regions, body sites manifest in zebrafish. To address this question, we sampled zebrafish guts at three locations (proximal, middle, and distal) after collecting feces from these animals. Since differences in microbial community composition across body sites (*e.g.*, gut vs skin) have also been well described in humans and other mammals ^44,46^ but not in zebrafish, we sampled three understudied non-gut sites (skin, gill, and swim bladder) in these same animals. Compared to non-gut sites (*i.e.*, skin, gill, and swim bladder), proximal gut (glmm; z = 3.28, *P* = 1.03x10^-3^), middle gut (z = 4.44, *P* = 8.91x10^-6^), and feces (z = 5.11, *P* = 3.23x10^-7^) had elevated richness (**Figure 2A**). A trend of increased richness in the distal gut microbiome was also observed but did not reach significance (z = 1.57, *P* = 1.11x10^-1^). In contrast, only the middle gut region had elevated Shannon entropy (z = 2.17, *P* = 3.00x10^-2^), indicating that differences in alpha-diversity between zebrafish tissue are primarily driven by an increased number of taxa not by increases in how evenly these taxa are distributed (**Figure 2B**).

**Figure 2.**
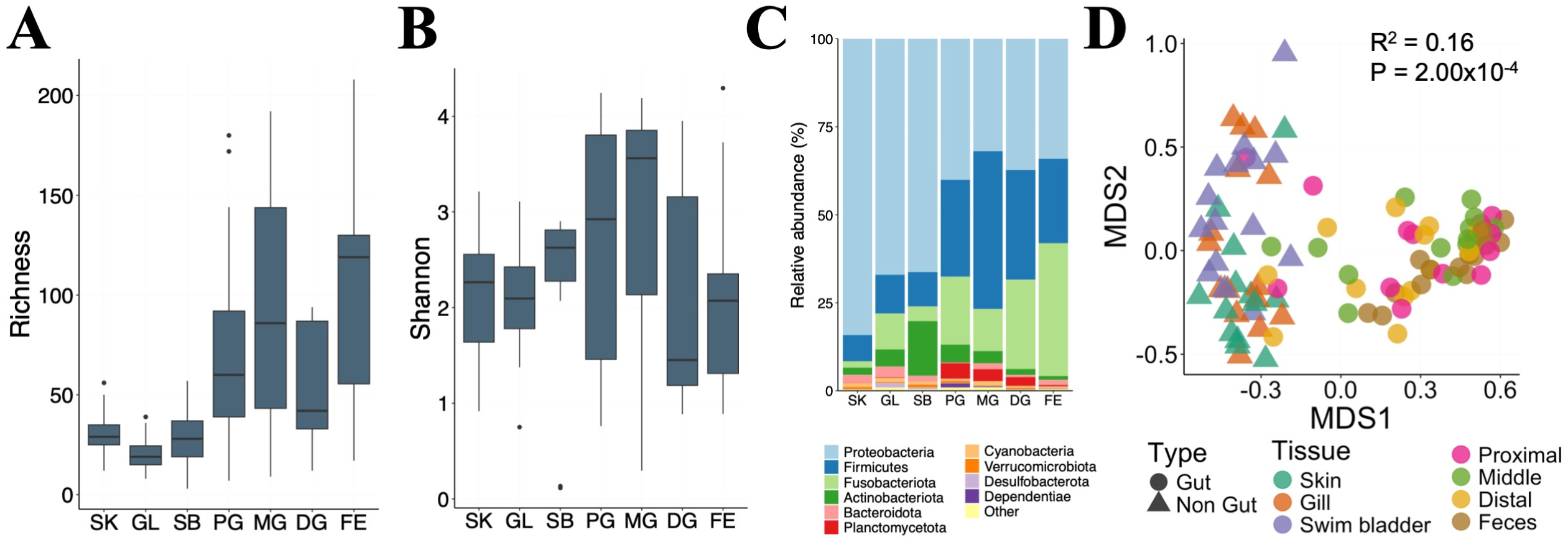
Zebrafish microbiome diversity varies significantly between tissues. **A)** Microbial richness and **B)** Shannon entropy at each body site sampled. **C)** Proportional abundance of the 10 most abundant phyla in each body site. **D)** An ordination of non-metric multidimensional scaling (NMDS) of microbial community diversity. The R^2^ value at the top left of the ordination indicates the strength of association between microbial beta-diversity and body site (PERMANOVA). SK: Skin, GL: Gill, SB: Swim bladder, PG: Proximal gut, MG: Middle gut, DG: Distal gut, FE: Feces.

Consistent with previous studies ^36,47–49^, we found that gut microbiomes of zebrafish were dominated by Proteobacteria, Fusobacteria, and Firmicutes (**Figure 2C, Supplemental Figure 2**). Proteobacteria and Firmicutes were the most abundant phyla for the gill, skin, and swim bladder. Despite sharing the same major phyla, microbial community diversity was significantly associated with tissue location (**Figure 2D**; PERMANOVA; R^2^ = 0.16, *P* = 2.00x10^-4^). Microbiome variation was also stratified by tissue type (*i.e.*, gut or non-gut), however, less variation was explained than by tissue (R^2^ = 0.07, *P* = 2.00x10^-4^). Since the PERMANOVA results indicated there may be tissue type specific microbial diversity in gut and non-gut tissues, we asked how taxa abundance varied between these two tissue types. A total of 32 genera significantly varied in abundance between gut and non-gut tissues (**Figure 3A**; GLMM, FDR < 0.05). Taxa with greater abundance in the gut included known zebrafish gut commensals *Cetobacterium* (z = -4.68, *P* = 3.35x10^-6^), ZOR0006 (z = -3.79, *P* = 1.49x10^-4^), and *Peptostreptococus* (z = -7.05, *P* = 1.81x10^-12^)^36^. Non-gut tissues had higher abundance of the genera *Shewanella* (z = 3.09, *P* = 2.03x10^-3^), and *Limnobacter* (z = 2.41, *P* = 1.58x10^-2^).

**Figure 3.**
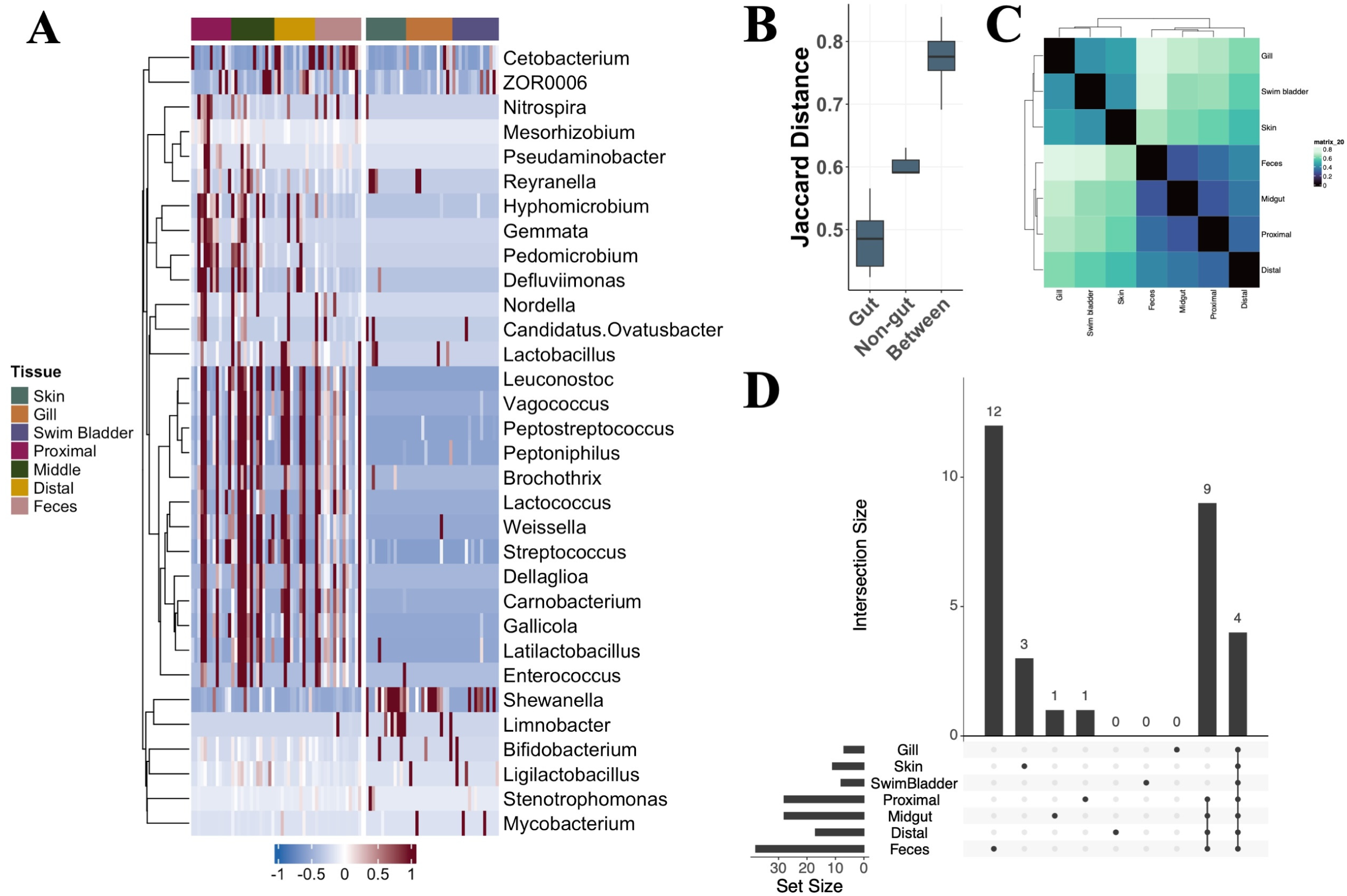
Zebrafish tissues possess substantial site specific and shared microbial biodiversity. **A)** A heat map of genera whose abundance varied across sampled tissues. Column color bar indicates tissue type. **B)** Jaccard distances within gut or non-gut tissues and between gut and non-gut tissues. **C)** A tile plot of Jaccard distance of genera prevalence between different tissues type. **D)** The upset plot shows the number of shared and unique genera (50% prevalence) between tissues (vertical bars) and the total number of genera identified in those tissues (horizontal bars).

To determine if tissues possess unique microbial signatures, we quantified the prevalence of each taxon in each tissue and then calculated the pairwise distance between all samples within and between groups. A taxon was considered present in a tissue if 50% or more of fish had a non-zero count for that taxon in that tissue. Overall, taxa prevalence in tissues of the same type (i.e., gut or non-gut) were more similar to one another that those in different tissue types (**Figure 3B, C**). Gut tissues were more similar to each other than were non-gut tissues (pairwise Wilcoxon test; W = 0, *P* = 0.02) and both gut (W = 0, *P* = 1.08x10^-4^) and non-gut (W = 0, *P* = 4.40x10^-3^) tissues were more similar to tissues of the same type than different tissue types (**Figure 3B**). Four genera *Cetobacterium*, *Aeromonas*, *Pseudomonas*, *Vibrio* were shared by all tissues (≥ 50% prevalence) regardless of type. Gut tissues uniquely shared an additional nine genera: *Peptoniphilus*, *Hyphomicrobium*, *Peptostreptococcus*, *Streptococcus*, *Weissella*, *Vagococcus*, *Gallicola*, *Leuconostoc*, and *Carnobacterium*. No genera or ASVs were uniquely shared between non-gut tissues at any prevalence threshold (**Figure 3C; Supplemental Figure 3**). Feces encoded the most unique diversity with 12 genera – *Candidatus Ovatusbacter, W5053, Stenotrophomonas, Enterococcus, Paucilactobacillus, Coxiella, Anaerosphaera, Lactobacillus, Erysipelothrix, Clostridium sensu stricto 5, Raoultibacter, Lachnospiraceae* AC2044 group – represented in only this compartment at 50% prevalence, followed by skin (*Limnobacter*, *Flavobacterium*, and *Sphingomonas*) with 3, then proximal gut (*Nitrospira*) and midgut (*Chloronema*) each with one (**Figure 3D**). Taken together these data suggest that gut and non-gut tissues are host to unique microbial communities.

### Gut and non-gut tissues possess unique functional potential

Since taxonomy varied across tissue types, we next asked if these differences resulted in distinct functional niches as well. Functional imputation suggests substantial differences in microbiome functional potential exists between zebrafish tissues (PREMANOVA; R2 = 0.13, *P* = 2x10^-4^). A total of 2,017 KEGG orthologous gene family abundances varied significantly across tissues (**Supplemental Table 3**; Kruskal-Wallis, FDR < 0.01). To determine if these alterations in individual gene families amounted to differential abundance of whole metabolic processes ecological niches in the zebrafish, we quantified differences in KEGG pathway abundances across tissues. Consistent with the gene family analysis, 140 KEGG pathways significantly varied between tissues (**Figure 4A**; **Supplemental Table 4**; Kruskal-Wallis, FDR < 0.01) many of which were involved secondary metabolism, xenobiotic biotransformation, and antimicrobial resistance, indicating that functional differences in zebrafish microbial niches extend beyond housekeeping genes.

**Figure 4.**
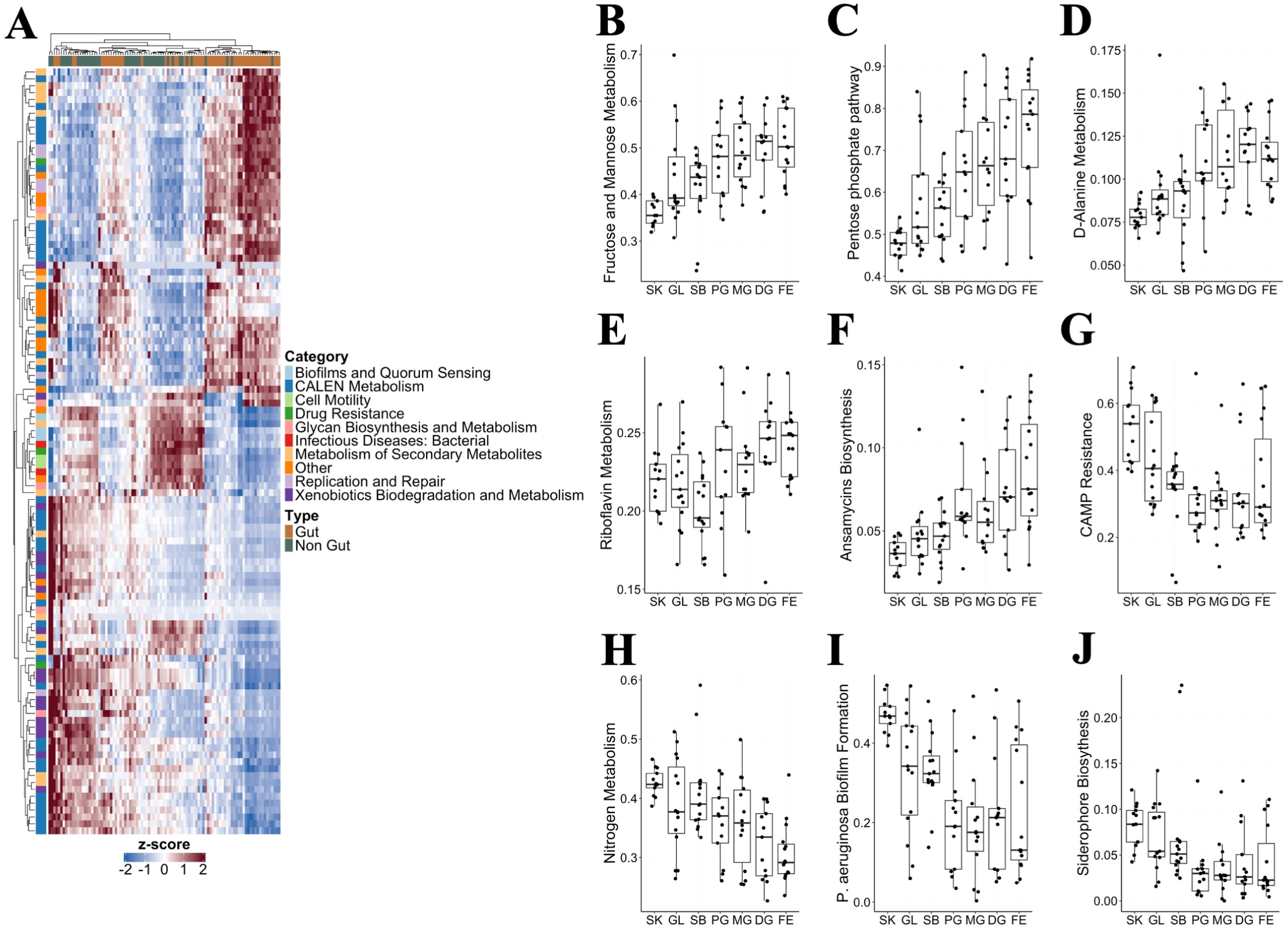
Microbial communities sourced from gut and non-gut tissues possess distinct functional profiles. **A)** a heat hap of scaled KEGG pathway abundances that significantly varied across sampling locations. Colored side annotations indicate the broad functional classification of pathways. Box plots show relative abundance of gene families which were increased in **B-F)** gut or **G-K)** non-gut tissues. SK: Skin, GL: Gill, SB: Swim bladder, PG: Proximal gut, MG: Middle gut, DG: Distal gut, FE: Feces.

**Figure 5.**
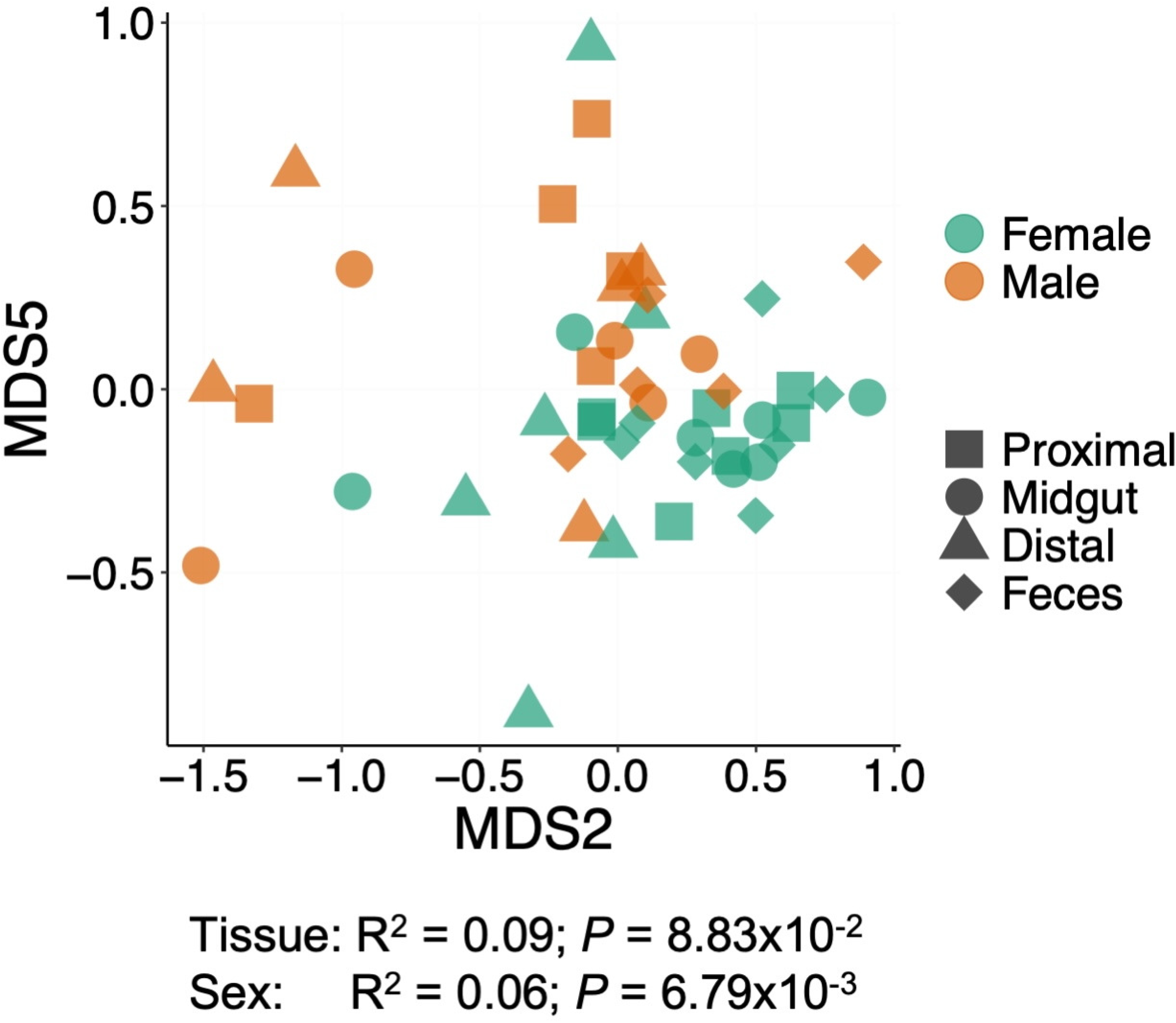
Zebrafish microbiome diversity associates with sex. Non-metric multidimensional scaling (NMDS) of gut tissue (shape) microbial community diversity in females (green) and males (orange). The R^2^ (PERMANOVA) and *P* value for associations between microbiome diversity, sex, and tissue are indicated below the ordination.

Functional category labels, based on KEGG BRITE categories, were assigned to each pathway to further classify these differentially abundant pathways according to broad functions. The most abundant functional categories identified were involved in Carbohydrate, Amino acid, Lipid, Energy, and nucleotide (CALEN) metabolism (41), xenobiotics biodegradation and metabolism (17), and in metabolism of secondary metabolites (17). Greater abundance of pathways associated with these categories was not restricted to one tissue type, but rather both gut and non-gut tissues had elevated abundance in specific members of the functional groups (**Figure 4A**). For example, gut tissues had higher abundance of pathways involved in metabolism of some carbohydrates (**Figure 4B-C**) including those generated during the break down of plant fiber. Gut tissues are also predicted to encoded higher abundance genes involved in D-alanine metabolism, a crucial step in peptidoglycan biosynthesis (**Figure 4D**). Pathways involved in production of antimicrobial compounds and production of B vitamins were also elevated in the gut (**Figure 4E-F**). In non-gut tissues notable examples of differentially abundant pathways included cationic antimicrobial peptide (CAMP) resistance, nitrogen metabolism, biofilm formation, and nutrient acquisition (**Figure 4G-J**). Together these results suggest that, in addition to distinct taxonomic composition, tissue sites in zebrafish likely encode unique functional potential.

### Sex shapes zebrafish microbial community diversity

A growing body of literature suggests that sex specific differences in gut microbiota link extensively with health and may potentially contribute to the pathogenesis of disease ^50,51^. However, little information is available on how sex may impact zebrafish gut microbial communities. In a third experiment we examined microbial community diversity in male and female zebrafish in feces and along the intestine. Neither richness (GLMM; *P* = 0.91) or Shannon entropy (*P* = 0.61) were impacted by sex in zebrafish gut microbial communities. However, a significant association between microbiome diversity and sex was identified (PERMANOVA; R^2^ = 0.06; *P* = 6.79x10^-3^). An association between gut compartment and diversity did not reach significance (R^2^ = 0.09; *P* = 8.83x10^-2^). However, while several genera were differentially abundant between tissue segments relatively few varied by sex (**Supplemental Table 5**; FDR < 0.2). The highly abundant genera *Aeromonas* (z = -2.41, *P* = 0.02) was lower in males, while the genera *Nocardioides* (z = 2.86, *P* = 4.04x10^-3^) showed higher abundance. Similar patterns of sex difference were observed at the ASV level. Several highly abundant gut taxa varied across intestinal location. For example, *Vibrio* abundance was higher in feces (z = 3.23, *P* = 1.00x10^-3^) while *Vagococcus* (z = -2.55, *P* = 1.06x10^-2^) and *Reyranella* (z = -3.52, *P* = 4.25x10^-4^) were decreased. In contrast, the genus *Reyranella* was increased in the proximal intestine (z = 5.98, *P* = 2.19x10^-9^) as was the genus *Coxiella* (z = 2.06, *P* = 2.19x10^-9^). The genus *Reyranella* was also increased in male zebrafish, however, this trend failed to reach significance (z = 1.88, *P* = 6.04x10^-2^). Together these data support a role for sex in shaping gut microbial community diversity in zebrafish.

## Discussion

As the use of zebrafish increases in microbiome research, there is growing need to develop guidelines and best practices for microbiome studies in this model. It is also critically important to define how microbiome ecology and operation in zebrafish corresponds to that of humans to identify application for this model translational relevance. The goal of this work was not to prescribe a universal set of husbandry parameters for microbiome studies in zebrafish, but rather to provide information on what factors influence experimental outcomes, and how these impacts may be mitigated or at least accounted for. Here we demonstrate that cohabitation, sex, and sampling location are critical determinants of microbiome composition and functional potential in zebrafish. These findings mirror those in other preclinical animal models and humans further arguing that, despite substantial taxonomic differences between zebrafish and humans, gut microbial communities operate very similarly in zebrafish and mammals^49,52–54^.

Cohabitation has been shown to have a homogenizing impact on the microbiota in mammals and can complicate interpretation of experimental results^43,55,56^. The impact of cohousing on microbiomes has been incompletely described in zebrafish. Our work has, for the first time, established that co-housing has similar homogenizing impacts on zebrafish microbiomes, and that these tank effects can be mitigated with individual housing. Unexpectedly, animals on recirculating flow systems have higher levels of richness and entropy and hosted more distinct microbiota which decreased significantly when placed on flow-through systems. Lower microbiome diversity in animals on flow-through could be due to incomplete sterilization of water on recirculating systems^34^ which in turn leads to stochastic and transient incorporation of a subset of these environmentally taxa into the gut microbiome. In this scenario, reduced biomass of water used on flow-through systems would result in a gradual reduction in microbial diversity, which is consistent with our observations. This gradual reduction in diversity subsided within 14 days of transfer to the flow-through system in both individually and group housed animals. Importantly, the rapid restructuring of the microbiome that occurs during equilibration likely could mask experimental impacts on microbial communities. Based on these findings an equilibration period of at least 14 days (individually housed) 21 days (group housed) is recommended prior to initiating experimental procedures when animals are transferred between tanks or systems.

In humans fecal microbial communities profiles imperfectly reflect intestine associated microbiota abundances^27^. This inconsistency can result in missing signatures of disease that can otherwise be detected by directly sampling intestinal tissue^37^. Here we demonstrate that zebrafish microbial community profiles also do not perfectly correlate with those of the intestine. This result is consistent with previous work in zebrafish larvae which demonstrated spatial heterogeneity in colonization dynamics along the intestine potentially due to competition or motility^57–59^. Alternatively, the intestine specific signatures may be due colonization microbes whose intestinal localization is due to specific nutrient requirements or interaction with spatially restricted gut cell types^60,61^. A limitation of this work was that we have used functional imputation which imperfectly models gene abundances using taxonomic signatures^41^. Shotgun metagenomics of intestinal and fecal samples would provide a less biased picture of microbiome functional potential which may better resolve potential mechanisms underlying these differential abundances. Regardless of the causes of compartmental differences in microbiome profiles in the gut our results argue for careful consideration of possible location specific effects when selecting a sampling strategy for zebrafish microbiome experiments. Whenever possible, it is recommended to collect intestinal samples at terminal experimental time points in case evaluation of intestine associated microbiota becomes necessary.

Diverse ecological niches have been described inside and outside the gut in humans and other animals^44,62,63^. While fecal communities are well explored in zebrafish ^36,47–49,64^, little is known about microbiome diversity at other sites. Consistent with previous work in zebrafish and other fishes, Proteobacteria, Fusobacteria, and Firmicutes dominated, gut, fecal, gill, and skin ^48,65–67^. The swim bladder – a gas filled sac connected to the gastrointestinal tract via the pneumatic duct which allows modulation of buoyancy – displayed similar patterns of microbial diversity. However, as in humans, even tissues close in proximity (*i.e.*, intestine and feces) displayed distinct patterns of diversity likely resulting location specific differences in nutrients, oxygen availability, chemical parameters, or host factors^37,44,68–70^. Our functional imputation analysis is consistent with this hypothesis as nutrient acquisition and metabolic pathways were the most often discriminatory between gut and non-gut tissues. For example, several pathways involved in vitamin production and utilization of simple carbohydrates were increased in the gut compared to non-gut tissues. This increase may reflect the greater levels of simple sugars available from the diet. At non-gut sites, particularly the skin and gill, nitrogen metabolism pathway abundance was elevated which may be due to the presence of low levels of ammonia, nitrate, and nitrite that are present in the environment^5^. Cationic antimicrobial peptide resistance pathway abundance was also increased in non-gut tissues. Such resistance strategies are likely necessary to for microbial colonization of as these sites as skin and gill tissues express high levels of CAMPs^71^. It is possible that some of the functional differences in gut and non-gut sites could reflect the increased exposure of non-gut sites to the environment (*i.e.*, water). A limitation of this work is that we did not sample the water column, so it is not possible to directly quantify the contribution of the water microbial structure at non-gut or gut sites.

However, a large study of marine fishes that tracked origin of gut, skin, and gill microbiota concluded that most taxa at each site were from an unknown origin and not water or sediment^72^. In zebrafish, others have found distinct microbial community structure in tank water and zebrafish gut^64^. Together these studies suggest that while exposure to the environment may contribute to microbial colonization in zebrafish, it is unlikely to be the primary driver of microbiome structure. Further shotgun metagenomic analyses are needed to clarify if and how the functional differences observed here may contribute to shaping microbial niches and host interactions in zebrafish.

Growing work demonstrates the role of sex and sex hormones in shaping the function and composition of microbial communities in humans and other mammals^73^. In zebrafish, sex specific differences in gastrointestinal gene expression have been reported^74^ as have sex dependent responses to perturbations on the microbiome^33,75^, however, conflicting reports exist on the existence of sex specific structure to zebrafish microbiomes. In this study we find that, consistent with mammals, sex does impact adult zebrafish, however these effects were relatively modest^73^. The reason for inconsistency in reports of sex differences in the microbiome of zebrafish is unclear, but could be related to husbandry effects (*e.g.*, flow, diet, housing density, etc.), technical limitations (*e.g*., primer bias), or other factors (*e.g.*, age, condition factor, etc.)^33^. It is also possible that prior work underpowered to detect subtle differences in the microbiome due to sex. The relatively small sample size used in this work also likely resulted in insufficient statistical power to resolve taxon specific differences between sexes resulting in an underestimation of sex specific microbiota differences. Given sex specific differences observed in other aquatic and terrestrial animals^73,76^ and recent reports that highlight sex dependent effects of diet on zebrafish microbiomes^22^ our data recommend inclusion of sex balanced designs in zebrafish microbiome studies.

Collectively, our findings demonstrate the substantial role that housing and sampling strategy play in shaping zebrafish microbiome composition. The current work was limited in scope and the diversity of factors that influence zebrafish gut microbiota far exceed what we were able to investigate. A community wide effort that includes facilities with diverse husbandry practices, zebrafish lines, and housing systems is needed to explore the factors underpinning microbiome composition and function and suggest standards of reporting and best practices for microbiome research in zebrafish. These efforts should incorporate functional as well as taxonomic measures of microbiome diversity, which would allow for a more detailed examination of how experimental parameters influence zebrafish microbial communities and how they relate to microbial communities of humans and other mammals. Finally, until standards and best practices can be developed, we recommended minimum reporting for zebrafish microbiome research include: feed type and feeding frequency; housing density; water quality parameters; housing system type; sampling strategy; light cycle; and fish line, age, sex, weight and length. These data would allow for greater reproducibility in zebrafish microbiome studies and potential provide insights into other factors that drive microbiome variation in zebrafish.

## Acknowledgements

This work was supported by an award from National Institutes of Health (R01ES036174) and institutional funds to CAG. We thank the University of Illinois Urbana-Champaign (UIUC) DNA Services Core at the Roy J. Carver Biotechnology Center for their assistance with sequencing. This work made use of the equipment, software, and facilities provided by the University of Illinois Urbana Champaign College of Veterinary Medicine Shared Equipment Program’s Biocomputing Shared Resource (BioShaRe). The College of Veterinary Medicine BioShaRe is housed in the Illinois Campus Cluster, a computing resource that is operated by the Illinois Campus Cluster Program (ICCP) in conjunction with the National Center for Supercomputing Applications (NCSA) and which is supported by funds from the University of Illinois at Urbana-Champaign.

---

**Supplemental Figure 1. Water flow type and housing strongly influence microbial community composition of zebrafish. A)** Experiential design and collection timeline. **B)** Longitudinal proportional genera abundance for group (top) and individually (bottom) housed animals.

**Supplemental Figure 2. Microbial community composition varies across tissue**. **A)** Phyla and **B)** genus bar charts show proportional abundance of the top 10 most highly abundant taxa in each microbiome. SK: Skin, GL: Gill, SB: Swim bladder, PG: Proximal gut, MG: Middle gut, DG: Distal gut, FE: Feces.

**Supplemental Figure 3. Zebrafish microbial communities share substantial microbial community diversity.** Upset plots showing **A-C)** ASVs or **D-E)** genera present in **A,D** at least 1 sample, **B)** 50% samples, or **C, E)** 80% samples shared or unique to a sampling site or set of sampling sites. Vertical bars show the number of taxa which overlap between each set of sites. Sets are indicated by connected vertical lines. Horizontal bars indicate the total number of ASVs or genera present at each sampling site.

